# Localized F-actin establishes critical cell interactions required for stem cell niche formation and identity

**DOI:** 10.64898/2026.08.13.744741

**Authors:** Rachael Johnson, Everette Rhymer, Robert Hughes, Lauren Anllo

## Abstract

Lifelong stem cells are maintained by a cellular microenvironment called the niche, which enables tissue homeostasis. Proper niche construction is essential for persistent function, but studying niche formation is challenged by the inaccessibility of most niches to *in vivo* visualization during development. Innovations imaging the *Drosophila* testis are now allowing investigation of niche inception. F-actin localizes to precise cell interfaces during testis niche assembly. Yet it is unknown whether this localization directs niche cell motility, or reflects adhesive sorting in response to formation of niche cell contacts. By adapting a method to optogenetically manipulate cortical F-actin via disruption of Rho1, we interrogate the role for cytoskeletal contributions to niche formation with tissue and temporal specificity. Rho1-mediated disruption of F-actin localization caused defects in niche anterior assembly and architecture. Also, fewer cells adopted bona fide niche identity, given diminished Fasciclin3, N-Cadherin, Islet, and Hedgehog. These disrupted niches fail in signaling to germ cells to establish stem cell identity. We reveal that localized F-actin is crucial for establishing cell contacts to form a functional niche, and to maintain cell identity in the developing tissue.

**Article Summary:** Stem cell niches are tissues that maintain lifelong stem cells, enabling regeneration. Constructing proper niche architecture during development is critical for tissue health and maintenance, but study of initial niche morphogenesis is challenged by niche inaccessibility to imaging and precise genetic manipulation. This study uses optogenetic cortical localization of cytoskeletal disruptors to reveal tissue and temporal specific requirements for localized F-actin in establishing a functional stem cell niche. This work unveils that F-actin driven formation of niche cell contacts during development is required to assume bona fide niche identity, which is essential to establish and maintain stem cells.

## Introduction

Stem cells are critical for homeostasis and regeneration of many adult tissues (Foster et al., 2002; Slack, 2018; Zakrzewski et al., 2019). To maintain a pool of non-differentiated stem cells required for tissue repair, stem cells require self-renewal signals from their cellular microenvironment, the niche. Niche cells often reside in specific tissue regions, restricting self-renewal signals to neighboring cells, thereby allowing cells out of signaling range to differentiate to maintain the tissue over time (Hicks and Pyle, 2023; Wagers, 2012). Stem cells are depleted causing tissue wasting when a niche does not develop properly, and the stem cell pool will over-proliferate generating tumors if there are excess self-renewal signals (Hicks and Pyle, 2023). Although niches have been identified in several adult tissues, mechanisms mediating their compartmentalization during development remain unclear.

The *Drosophila* testis is a paradigmatic model revealing concepts in stem cell niche biology, as it has a well-defined niche, is genetically tractable, and is accessible to *in vivo* live imaging (de Cuevas and Matunis, 2011; Hales et al., 2015; Nelson et al., 2020). The adult testis is a coiled tube with the spermatogonial niche compartmentalized at the tip (Hardy et al., 1979). Germline stem cells (GSCs) radially surround the niche and divide perpendicular to the niche-GSC interface, ensuring that GSC daughter cells are displaced away from the niche environment and will differentiate, eventually generating sperm (Yamashita et al., 2003). Niche signals including Bone Morphogenetic Protein (BMP), and those acting through Janus kinase signal transducer and activator of transcription proteins (JAK-STAT) enable germline stem cells to maintain fertility throughout adulthood (Leatherman and DiNardo, 2010; Tulina and Matunis, 2001).

The testis niche is established during late gonadogenesis in embryonic stages (ES) 15-17 (Anllo et al., 2019; Le Bras and Van Doren, 2006). Gonad formation requires coalescence of somatic gonadal precursors (SGPs) with germ cells. Prior to coalescence, SGP receipt of Notch signaling is required for specification of niche cell fate (Okegbe and DiNardo, 2011). Later in the coalesced gonad, a subset of embryonic SGPs assembles a spherical niche structure at the embryonic gonad anterior (Kitadate & Kobayashi, 2010; Le Bras & Van Doren, 2006; Okegbe & DiNardo, 2011). Some pro-niche SGPs are localized at the gonad anterior at the onset of gonad coalescence, while others must migrate onto extracellular matrix at the gonad periphery and move towards the anterior to assemble the final niche. Additional SGPs remain interspersed among and encysting germ cells in more posterior regions of the gonad after niche assembly (Anllo et al., 2019). When pro-niche SGPs cluster, F-actin is enriched first at niche-niche cell interfaces, and later re-localized to niche-GSC interfaces (Fig. 1A) (Anllo and DiNardo, 2022). It remains unclear whether F-actin localization directly mediates cell motility required for niche assembly, or whether it reflects adhesive sorting in response to formation of niche cell contacts. Upon genetic disruption to assembly cues, niche cells retain accumulation of niche-specific adhesion proteins despite their dispersed, mislocalized position, suggesting that adhesive sorting alone is not sufficient for proper niche morphogenesis. Yet a direct role for cortical F-actin enabling assembly has not been tested, and cytoskeletal regulators mediating F-actin localization during niche assembly are unknown.

**Figure 1.**
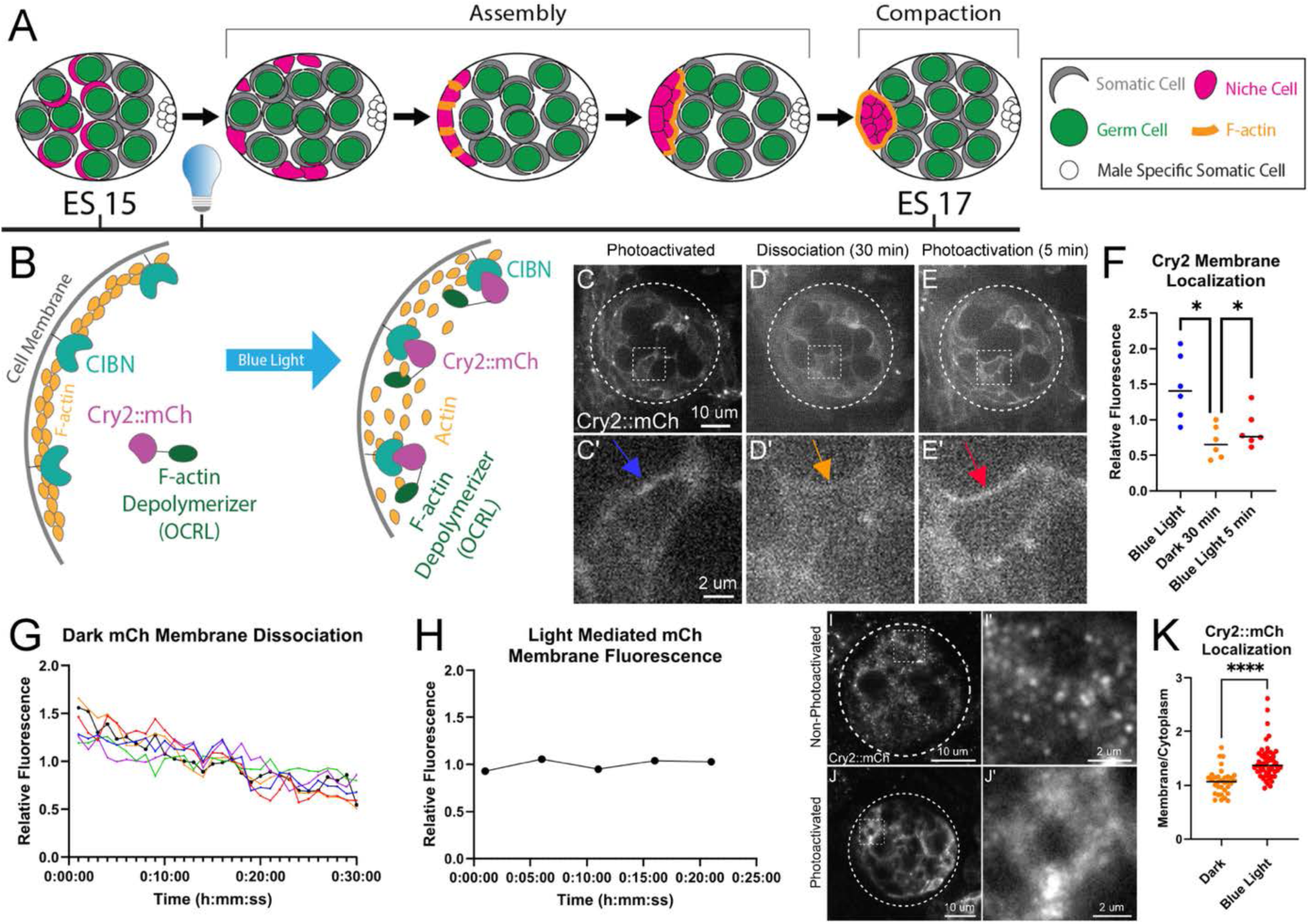
Optogenetic methods mobilize F-actin modulators to SGP cortices. (A) Diagram showing F-actin localization during niche assembly. In experiments, blue light is applied after embryonic stage 15 begins and just prior to niche assembly (light bulb). (B) Blue light induces oligomerization of Cry2-tethered proteins with membrane-tethered CIBN. (C-E) Live imaging of Cry2-mCherry showing cortical localization after blue light photoactivation (C), followed by reduced cortical accumulation after 30-minutes of dark (D), then re-accumulation at the cortex upon blue light reapplication (E). Prime panels represent outlined boxes showing one enlarged cell at each timepoint. (F) Quantified Cry2-mCh cortical enrichment in photoactivated vs dark dissociated cells (p = 0.0313, n = 6 from 3 gonads, Wilcoxon test), and dark vs re-photoactivated cells (p = 0.0312, Wilcoxon test). (G) Negative correlation of Cry2-mCh cortical fluorescence over 30 minutes without blue light (p<0.0001, r = -0.9739, Pearson correlation, n = 6 cells from 3 gonads). (H) Average cortical Cry2-mCh during photoactivation (n = 4 cells). (I, J) Immunostained Cry2-mCherry localization in dark-state non-photoactivated (I, I’) and photoactivated tissue (J, J’). (K) Cortical Cry2-mCherry enrichment in non-photoactivated (Dark) and photoactivated (Blue light) embryos (p<0.0001; Dark, n = 33 SGPs from 9 gonads; Blue light, n = 56 SGPs from 15 gonads; Mann-Whitney test). n ≥ 2 trials. **Alt Text:** Diagrams illustrating F-actin enrichment during testis niche assembly, and cortical recruitment of optogenetic constructs upon blue light application. Accompanying images of cells highlight cortical enrichment of Cry2-mCh modulated by light exposure, which is quantified in several associated graphs with statistical annotations.

Assembled niche cells rearrange to form the final, compact niche architecture after assembly and later in embryogenesis. This final structure is maintained in the functional adult niche. Rho kinase mediated actomyosin contractility compacts assembled niche tissue into a spherical structure that restricts delivery of niche signals to a limited pool of germline stem cells (GSCs). At this later developmental timepoint, Rho kinase activity is not essential for niche cells to retain their identity or signal production, but is required to maintain the compacted niche shape that restricts the GSC pool during both embryogenesis and adulthood (Vida et al., 2025; Warder et al., 2024).

Testing the role for F-actin localization during initial niche assembly was previously hampered by the earlier requirement for F-actin regulation in earlier gonad coalescence (Sano et al., 2012). To perturb F-actin localization during niche assembly, we adapted the Cry2-CIB optogenetic system previously developed to study embryonic epithelial morphogenesis (Guglielmi and De Renzis, 2017). Using this approach, we disrupt SGP F-actin precisely during the developmental timepoint of niche assembly. We show that F-actin is localized by somatic Rho1, and this localization is required to establish cell contacts required for incipient niche assembly. We find Rho1 dependent F-actin localization, mediated in part by Ena/Vasp, is required for accumulation of niche cell adhesion proteins and expression of the niche identity markers Islet and Hedgehog. This work identifies a critical role for F-actin localization in both establishing niche cell contacts and niche cell identity during the assembly process. These data suggest that cytoskeletal regulation is a key mediator of niche formation and function to maintain tissue homeostasis over the lifetime of the organism.

## Materials and Methods

### Experimental methods

#### Drosophila stocks

All *Drosophila* lines are listed in the key resource table and genotype tables (Table 1 and Supplemental Tables). For optogenetic experiments, UAS-Cry2-CIBN fly lines were crossed to *Six4*-Gal4::VP16 genetic driver lines containing a *Six4-*eGFP::moesin sequence to generate embryos in which we could visualize somatic F-actin live. For optogenetic experiments with fixed tissue analysis, UAS-Cry2-CIBN fly lines were crossed to *Six4*-Gal4::VP16 genetic driver lines, and embryos were immunostained for a fluorescent gonad marker.

**Table 1.** Genotype Table.

| **Experiment** | **Light-exposed Control Genotype** | **Manipulated and Dark State Genotype** |
|---|---|---|
| OCRL Optogenetics | UAS-CIBN::CAAX ; CyO Dfd-GFP ; UAS-Cry2::mCh::OCRL | UAS-CIBN::CAAX ; *Six4*-Gal4::VP16 ; UAS-Cry2::mCh::OCRL |
| Rho DN Optogenetics | UAS-CIBN::CAAX ; CyO Dfd-GFP ; UAS-Cry2::mCh::RhoDN | UAS-CIBN::CAAX ; *Six4*-Gal4::VP16 ; UAS-Cry2::mCh::RhoDN |
| OCRL Optogenetics with F-actin label | UAS-CIBN::CAAX ; CyO DfdGFP ; UAS-Cry2::mCh::OCRL/*Six4*-eGFP::moesin | UAS-CIBN::CAAX ; *Six4*-Gal4::VP16; UAS-Cry2::mCh::OCRL/*six4*-eGFP::moesin |
| RhoDN Optogenetics with F-actin label | UAS-CIBN::CAAX ; CyO DfdGFP ; UAS-Cry2::mCh::RhoDN/*Six4*-eGFP::moesin | UAS-CIBN::CAAX ; *Six4*-Gal4::VP16 ; UAS-Cry2::mCh::RhoDN/*Six4*-eGFP::moesin |

#### Sex identification and genotyping

Male gonads were identified by observing male-specific SGPs (msSGPs) located at the posterior of the gonad, labeled with either a Vasa antibody which labels both germ cells and msSGPs, or *Six4*-eGFP::moesin which labels msSGPs with more robust fluorescence than other SGPs (Renault et al.; 2012; Sano et al., 2012). Sibling control embryos were identified using a fluorescently marked balancer (CyO Dfd-GFP). For UAS-Cry2::OCRL::mCh fly lines, genotypes were determined by observing red mCherry fluorescence.

#### Optogenetic photoactivation

All embryos and adults from optogenetic crosses were kept in dark boxes and processed using amber-filtered lights to prevent premature photoactivation. For live analysis experiments, embryos expressing Cry2::mCh and CIBN::CAAX in SGPs were photoactivated with a 488nm laser, then immediately imaged using a 555nm laser to visualize Cry2::mCh. Images were captured approximately every 5 minutes using a 35-60 μm Z stack with 0.5 μm intervals. Time intervals varied between 3.5 to 7 minutes to maintain photoactivation of optogenetically manipulated embryos. Live imaging experiments maintained photoactivation with an LDI 488 nm laser used to image GFP through a Crest X-Light V3 spinning disk confocal unit at 20% power and 300 ms exposure time with a 60x 1.3 NA silicone immersion objective. The interval of image capture was set to continuously image once cycled through positions (approximately 5 minutes) to maintain photoactivation throughout imaging sessions for 4 hours.

Fixed analysis experiments used a blue LED pulsed lightbox with a 300 ms exposure time to maintain photoactivation. Experiments maintained photoactivation in live, dechorionated embryos kept wet in a nytex collection basket from embryonic stage 15 to 17, which consisted of a 4-6-hour photoactivation period prior to dissection and fixation. Experiments to test Cry2 mobilization to the cortex were photoactivated for 4 hours, while all functional assessments of niche assembly utilized 6 hours of photoactivation.

#### Optogenetic genetic and dark-state controls

All optogenetic experiments were paired with genetic controls, which lacked the Gal4 driver but were exposed to our blue light protocol. Fixed tissue experiments were done with both genetic controls, and an additional set of dark-state controls, which had all optogenetic constructs but were reared completely in the dark. Fixed dark-state experiments used dark boxes to collect and age embryos that were dissected under amber filtered lights until fixation with 4% paraformaldehyde.

#### in vivo live imaging

Embryos were prepared for live imaging as described previously (Ong et al., 2019). Embryos were collected overnight (∼16 hours) at 25°C on an agar plate and dechorionated the next morning in a nytex basket using 50% bleach. Embryos were then selected for embryonic stage 15 using gut morphology to determine age according to Campos Ortega and Hartenstein (Campos-Ortega and Hartenstein, 1997). Embryos were mounted dorsolaterally to a slide using double-sided tape dissolved in heptane to create an adhesive solution and then covered in halocarbon 27 oil. Two 18x18mm coverslips were glued to the slide on both sides of the embryos to create a bridge. A 22x30 mm coverslip was then glued to the bridge coverslips to allow space for the embryos. Slides were imaged using an Olympus IX83 microscope with a Crest X-Light V3 spinning disk, using a 60x oil immersion objective.

#### Embryonic gonad dissection and immunostaining

Embryos were collected on grape agar plates for two hours at 25°C and aged for 22-24 hours at 18°C until embryonic stage 15, just prior to niche assembly. Collection and aging was done in a dark environment using amber filter lights and dark boxes to prevent premature photoactivation. Once aged to embryonic stage 15, embryos were dechorionated in a nytex basket with 50% bleach, and then photoactivated using a blue LED light pulsing with 300 ms exposure. Photoactivation was performed for 6 hours at 25°C, until the end of niche assembly at embryonic stage 17, followed by dissection and fixation in 4% paraformaldehyde (PFA) for 15 minutes for all experiments except STAT, which were fixed in 8% PFA for 30 min at 4C. All tissue was blocked using 4% normal donkey serum (NDS). Primary antibody staining was performed overnight at 4°C for all antibodies except STAT, which was done at room temperature. All secondary antibodies were used at 3.75 μg/ml (Alexa 488, Cy3, Alexa 647; Molecular Probes; Jackson Immunoresearch) for 2 hours at room temperature. Hoechst 3342 (Sigma) at 0.2 μg/ml for 5 minutes was used to stain DNA. All washes were done using phosphate-buffered saline with 0.1% Triton (PBST).

Rabbit antibody was used against RFP at 1:500 (abCam 62341), chicken antibody against GFP at 1:1000 (Cat#GFP-1020), goat antibody against vasa at 1:800 (Cat# sc-26877 dC-13), rabbit antibody against vasa at 1:5000 (Boster Bio DZ41154), mouse antibody against mCherry at 1:30 (DSHB-3A11 AB-2617430), rat antibody against N-cadherin at 1:50 (DSHB-DN-Ex #8 AB-528121), mouse antibody against Fasciclin3 at 1:20 (DSHB 7G10), rabbit anti STAT92E 1:200 (gift from E. Bach, Flaherty et al., 2010), mouse anti Islet 1:200 (DSHB AB_528313), and mouse anti Gamma Tubulin 1:200 (Sigma AB_528441). The phosphatase inhibitor, PhosStop (Sigma – Cat#4906845001), was used during dissections and immunostaining staining for experiments with STAT antibody.

Fixed tissue was imaged either with the Crest X-Light V3 spinning disk confocal (see above) or with a Zeiss AxioImager.Z1 equipped with ApoTome.2 and 40x 1.2 NA water immersion objective.

#### Niche phenotypic characterization

Normal niches were identified in live movies using the somatic cell marker *Six4*-eGFP::moesin as a compact and smooth cluster of cells at the gonad anterior, directly opposite of the msSGPs. Both niche cells and msSGPs express *Six4* at higher levels than other somatic gonadal cells, allowing for the identification of each. Normal niches in fixed tissue experiments were identified by immunostaining for the niche-specific adhesion protein markers N-cadherin and Fasciclin3. Abnormal niches were identified using *Six4-*eGFP::moesin, Fasciclin3, and N-cadherin and characterized by having irregular boundaries and/or not positioned at the anterior.

#### Visualization of normalized F-actin and Cry2::mCh fluorescence

F-actin in live movies was visualized by transgenic expression of *Six4*-eGFP::moesinGFP. Cry2::mCh in live movies was visualized by observing the mCherry tagged fluorophore. Cry2::mCh was visualized in fixed tissue by immunostaining for the mCh tag.

### Statistical Analyses

#### Quantifications

All fluorescence accumulation measurements were completed using ImageJ to report mean gray values for selected regions of interest (ROIs) at a z slice in which the interface was best in focus. All measurements were first background subtracted and then normalized as described for each experiment below.

#### Measuring F-actin at niche cell interfaces

Interface fluorescence measurements were taken by tracing niche cell-niche cell or niche cell-germ cell interfaces. Background fluorescence was measured as mean gray values at germ cell centers, since germ cells do not express *Six4*. Background subtracted values were normalized, dividing each of them by the average of all background subtracted values for their respective gonad.

#### Cry2::mCh recruitment and dissociation experiments

Fluorescence was measured around the entire cell cortex at every 1-minute time point for 30 minutes. Background fluorescence was measured as the value for a neighboring germ cell center at each time point. Each measurement was background subtracted at each timepoint. Background-subtracted measurements were normalized individually for each cell, dividing the measurement at each timepoint by the average value for that cell over the 30 min experimental period. Quantification of Cry2::mCh membrane recruitment in *Six4*-Gal4::VP16 manipulated embryos was normalized by dividing each background subtracted measurement by the average of all values for the respective gonad.

#### Cry2::mCh dark versus light state fixed experiments

ROIs were taken around the entire cell cortex and around the nuclei for relative fluorescence of membrane-to-cytoplasm ratios. These measurements were taken blindly using the somatic F-actin marker *Six4*-eGFP::moesin to delineate the cell cortex and Hoechst DNA stain to indicate nuclei. Background fluorescence was measured at a neighboring germ cell center. Background-subtracted measurements were normalized, dividing each of them by the average of all background-subtracted values for their respective gonad.

#### Measuring STAT accumulation

Gonads were stained with a Stat antibody (E. Bach; 1:400). ImageJ was used to measure the mean value fluorescence intensity. Regions of interest (ROIs) were selected from 5 GSCs and 3 neighboring germ cells, using solely Vasa as a germ cell marker and one Z plane with the germ cell best in focus. The fluorescence intensities of each germ cell were background subtracted, using background measurements from ROIs with no tissue. The ratio of Stat accumulation for each GSC normalized relative to the neighboring germ cell average for each individual gonad was measured. The relative Stat enrichment values for each GSC was plotted for control and mutant genotypes for RhoDN and *ena* RNAi. Mann-Whitney tests were used to compare the differences between the genotypes.

#### Measuring Ena/VASP at niche cell interfaces

Interface fluorescence measurements were taken by tracing niche cell-niche cell or niche cell-germ cell interfaces. Background fluorescence was measured as mean gray values outside of the gonad with no tissue. Background-subtracted values were normalized, dividing each of them by the average of all background-subtracted values for their respective gonad.

#### Measuring Hedgehog accumulation

Mean gray value fluorescence measurements were taken for 3 niche cortices where Ncad was detectable, with interfaces belonging to 3 different cells per gonad quantified. The segmented line plugin was used for ROI selection in ImageJ. For measurements from each gonad, background fluorescence from where no tissue was present was subtracted from mean gray values, which were then normalized to background-subtracted values taken from the periphery of a germ cell distant from the hub from the same gonad.

#### Measuring Islet accumulation

Mean gray value fluorescence measurements were taken for 3 niche cell regions within N-cad labeled niche cortical boundaries using the oval function in ImageJ for ROI selection. To quantify background fluorescence, an ROI was selected from a region where no tissue was present. Background-subtracted fluorescence values for each niche cell were normalized, dividing by the background-subtracted value from a neighboring unlabeled germ cell from the same gonad. Relative fluorescence values for each niche cell were plotted.

#### Measuring niche cell counts

Quantification of niche cells expressing niche markers, including N-cad or Fas3, was done using ImageJ Cell Counter Plugin. Embryonic niche cells accumulate Fas3 at contacts with other niche cells, and accumulate N-cad on all interfaces. Niche cells that express these niche specific adhesion proteins are co-labeled with Hoechst as a nuclear marker for distinction of cells.

All statistical analysis was performed using GraphPad Prism version 10 or newer. For all statistical analysis, significant comparisons are shown as follows: ns = note significant, *p value < 0.05, **p value < 0.01, ***p value < 0.001, and ****p value < 0.0001. All graphs display the significant comparisons, with p values noted in figure legends. Niche phenotypic characterizations were quantified using Fisher’s Exact test. Statistical comparisons of fluorescence intensity were evaluated with Mann-Whitney tests.

## Results

### Cry2-CIBN optogenetic system recruits F-actin modulators to niche cell cortices

SGPs dynamically encyst germ cells to form the nascent gonad. Thus, early somatic disruption to the F-actin cytoskeleton precludes a targeted interrogation of the role for F-actin localization specifically during gonad niche assembly. To address this, we employed Cry2-CIB optogenetics for precise temporal control to disrupt F-actin after the gonad first forms, specifically during the period of niche assembly. Cry2 and CIB are proteins derived from *Arabidopsis* that minimally interact in the dark and oligomerize in the presence of blue light. This interaction has been harnessed to enable light-mediated subcellular protein re-localization immediately upon blue light introduction (Kennedy et al., 2010). We used UAS controlled tissue-specific expression of Cry2 fusions for either of two cytoskeletal disruptors: the actin depolymerizer inositol-polyphosphate-5-phosphate OCRL (Cry2::OCRL::mCherry) or dominant-negative Rho1 GTPase (Cry2::RhoDN::mCherry) (see Table 1 and Supplemental Tables for all resources and genotypes) (Guglielmi and De Renzis, 2017; Guo et al., 2022). The mCherry tag allows visualization of sub-cellular localization. In embryos also expressing CIB tethered to the cortex (CIBN::CAAX), blue light will activate Cry2-CIB binding, localizing the cytoskeletal disruptors to cortical locations at or near the site of F-actin polymerization (Fig. 1B).

Previous applications used this system in superficially located ectoderm (Guglielmi and De Renzis, 2017), so we needed to validate its effectiveness in the internally positioned gonad. We used the SGP-specific *Six4*-Gal4::VP16 driver (Warder et al., 2024), and measured SGP cortical Cry2::OCRL::mCh localization in living embryos. Live imaging niche assembly *in vivo* requires precise orientation of embryos using a fluorescent marker to delineate the incipient gonad (Anllo et al., 2019). We used *Six4*-eGFP::moesin embryos for *in vivo* imaging somatic gonadal F-actin, leaving the red channel free for detection of membrane localized mCh. Embryo alignment using the *Six4*-eGFP::moesin marker during tissue preparation required blue light exposure and served as initial live photoactivation for embryos reared in the absence of blue light and aged precisely to the timepoint of niche assembly. After orienting embryos for optimal imaging, these were mounted on slides and live imaged with exposure to 555 nm light excitation to visualize mCh, and 488 nm excitation both to visualize *Six4*-eGFP::moesin and photoactivate. In photoactivated tissue, we observed Cry2::mCh enrichment at SGP cortices, suggesting response to blue light exposure (Fig. 1C,C’). To test the dissociation of Cry2 cortical localization in the absence of blue light, we continued live imaging for 30 minutes using only a 555nm laser. We quantified relative accumulation of Cry2-mCh at the cortex compared to the cytoplasm for individual SGPs selected because they remained visible at each minute for the entire 30-minute sampling period. We tracked two such cells per gonad for three trials with separate animals, for a total of 6 cells (Fig 1D,D’,G). We observed a negative correlation of mCh cortical enrichment over time (Fig 1G), representing progressive cortical reduction of Cry2::mCh fluorescence.

Comparison of cortical enrichment at timepoints 0 and 30 min showed a total reduction in cortical mCh accumulation (Fig. 1F). The progressive and total cortical reduction in mCh accumulation represent Cry2-mCh membrane dissociation when blue light is removed. This indicates that continued blue light exposure is required to maintain cortical recruitment. To confirm the progressive mCh reduction was dissociation and not an artifact of photobleaching, in other embryos we measured stable Cry2::mCh at the cell cortex over 30 minutes with continual photoactivation (Fig. 1H). After dissociation, we tested Cry2 recruitment back to the cortex, reintroducing 488nm blue light. We detected Cry2::mCh re-accumulation at the membrane in each of the 6 cells in which we tracked dissociation immediately (Fig. 1E&F). These results confirm light responsive mobilization of optogenetic constructs at SGP cortices during live imaging.

We would need to examine certain functional and gene expression consequences of optogenetic manipulations to niche assembly using immunofluorescence. Therefore, we also measured Cry2::mCh in embryos that were precisely aged to the developmental timepoint when niche assembly begins in the dark, then dechorionated and photoactivated with a pulsing blue LED light for 4 hours prior to dissection and fixation. Because light-independent Cry2-CIB interactions can occur (Kennedy et al., 2010), we compared Cry2::mCh membrane accumulation in non-photoactivated embryos (Fig. 1I), and genetically identical embryos photoactivated for 4 hours with blue light before processing (Fig. 1J). In these experiments, SGP cortices were delineated by *Six4*-eGFP::moesin. Results revealed significantly increased membrane-to-cytoplasm fluorescence after photoactivation in the experimental group (Fig. 1K), indicative of Cry2::mCh cortical recruitment. In non-photoactivated (dark state) embryos, low Cry2::mCh membrane recruitment suggests low-level dark state activity (Fig. 1K) indicating minimal light-independent interactions. Together, our data present the opportunity to use blue light exposure for precise temporal localization of cytoskeletal disruptors to the SGP cortex in both live and fixed tissues.

### Cytoskeletal modulator Rho1 is required for niche F-actin localization to enable niche assembly

Our previous work revealed F-actin localized at precise subcellular interfaces in assembling niche cells, initially enriched at niche-niche contacts and later re-localized to niche-stem cell interfaces (Anllo and DiNardo, 2022). During morphogenesis, localized F-actin at discreet subcellular interfaces can direct cell motility, or it can reflect adhesive sorting in response to establishment of cell contacts (Tsai et al., 2022). To test whether F-actin localization was required to direct niche formation, we used Cry2-CIB optogenetics to perturb SGP F-actin at the precise developmental timepoint when niche assembly occurs. Disruptions to assembly would suggest an active role for F-actin directing early morphogenesis, while unperturbed assembly would suggest that F-actin is localized in consequence of niche cell contact.

We first tested whether cortical F-actin was required for niche assembly. We optogenetically localized cortical OCRL to depolymerize F-actin during the period of assembly (Fig. 1A) and monitored results live in real time. Manipulated gonads inherited the *Six4*VP16-Gal4 genetic driver along with UAS-driven optogenetic components, while control animals did not inherit the *Six4*VP16-Gal4 driver (See Table 1 and Supplemental Table 2 for genotypes). In manipulated gonads, we detected Cry2::OCRL::mCh at SGP membranes that was absent from sibling control embryos lacking the *Six4*VP16-Gal4 driver (Fig. 2A’,B’). Control SGP membranes were delineated with *Six4*-eGFP::moesin to label somatic F-actin (Fig.2A,A’’). After 4 hours of *in vivo* imaging, sibling control embryos had enriched somatic F-actin in a clearly assembled anterior niche structure (Fig. 2A’’). In stark contrast, optogenetically manipulated animals showed punctate somatic F-actin throughout the gonad (Fig. 2B’’) and a corresponding failure to establish a singular anterior niche tissue (Fig. 2B’’-C). These results indicate a requirement for cortical F-actin during niche assembly. To confirm our live analysis results, we also photoactivated embryos of the same genotypes with pulsing blue LED light for 6 hours during niche assembly prior to dissection, fixation, and immunostaining. Fixed tissue analysis revealed assembled niches in controls lacking the *Six4*VP16-Gal4 genetic driver, and failed assembly with dispersed, non-anterior niche cells in manipulated gonads (Fig 2D-F). Because our experiments restricted blue light exposure to the assembly period, they bypassed the earlier requirement for F-actin during gonad coalescence. Our results reveal a temporal requirement for cortical F-actin during the niche assembly stage.

**Figure 2.**
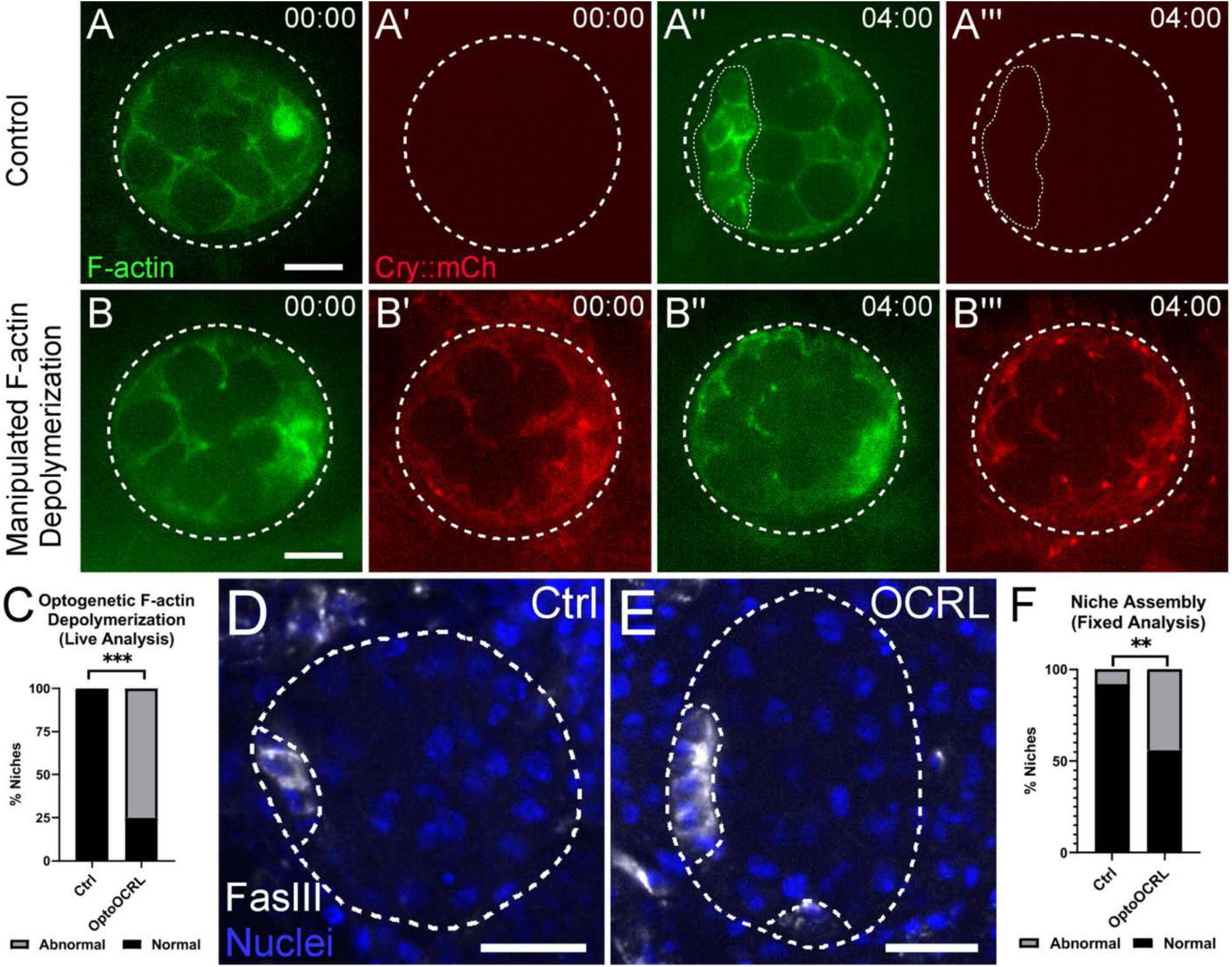
Cortical F-actin is required for niche assembly. (A,B) Live imaging control (A) and manipulated (B) gonads. Somatic F-actin, green. Cry-OCRL-mCh, red. Control niche outlined. (C) Quantified niche assembly frequency in controls and optogenetic OCRL-manipulated SGPs (p = 0.0003; Ctrl, n = 12 gonads; OCRL, n = 12 gonads; Fisher’s Exact test). (D,E) Control and optogenetic OCRL manipulated gonads (niche boundaries outlined: Fas3, white; Nuclei, Hoechst, blue). (F) Frequency of assembly defects with optogenetic OCRL manipulated gonads (p = 0.0084; Ctrl, n = 19 gonads; OCRL, n = 10 gonads; Fisher’s Exact test). Gonads outlined. Scale bars, 10 um. n ≥ 3 trials. **Alt Text:** Microscopy images of embryonic gonads show cortical localization of Cry2-OCRL-mCh in light-manipulated tissue, with failed assembly of the stem cell niche. Control images without cortical mCh fluorescence and normal niche assembly are depicted for comparison. Graphs with statistical annotations show that niches are abnormal with higher frequency in optogenetically manipulated tissue.

Actomyosin contractility modulated by Rho Kinase (Rok) defines and maintains assembled niche tissue architecture (Warder et al., 2024; Vida et al., 2025), but a role for Rok in initial assembly of niche cells has not been described. We tested a role for the Rok regulator Rho1 in initial niche assembly. We used live imaging with *Six4-*eGFP::moesin to visualize somatic F-actin in optogenetically manipulated gonads (Cry2::RhoDN::mCherry) (Fig. 3A-D). For individual niche cells, we quantified F-actin accumulation at interfaces with other niche cells, and interfaces with germ cells when niche cells began anterior clustering (00:00, hh:mm) (Fig. 3A’,C’) and after 1.3 hours (Fig. 3B’,D’). In controls, as expected from our prior work, we observed enriched F-actin at niche-niche interfaces when niche contacts were first established (Fig. 3A’), switching to niche-germ cell interfaces after 1.3 hours (Fig. 3B’). However, in photoactivated gonads with RhoDN recruited to the cortex, F-actin was still cortically enriched but was never polarized (Fig. 3C-D’), revealing that Rho1 is necessary for subcellular localization of F-actin in niche cells during assembly.

**Figure 3.**
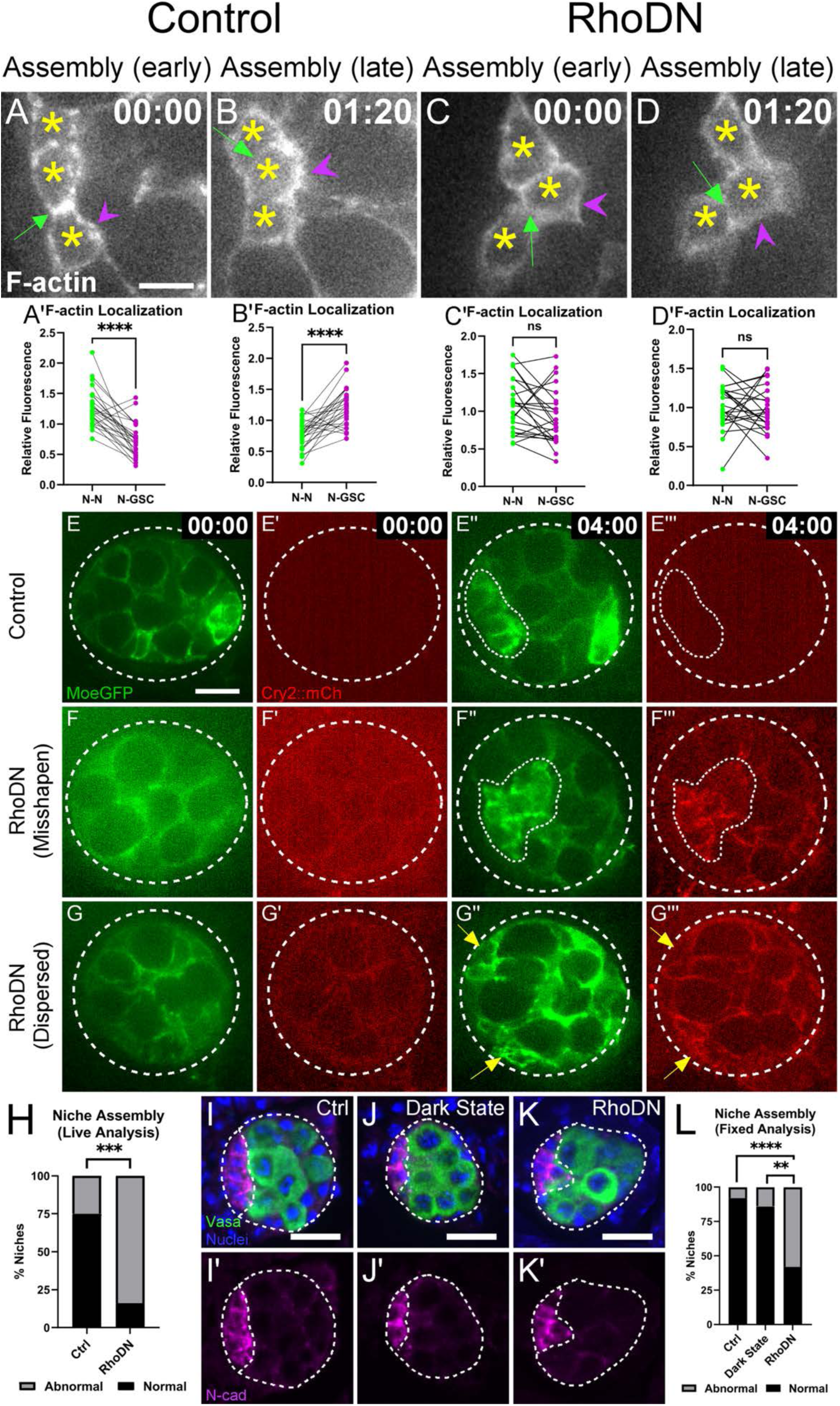
Rho1 mediated F-actin localization is required for niche assembly. (A-D) Live imaging F-actin at niche-niche (green) and niche-GSC (magenta) interfaces, quantified in prime panels. Niche cells, asterisks. (A’,B’) p<0.0001 (n = 25 cells from 10 gonads). (C’) p = 0.0564. (D’) p = 0.4908 (n = 24 cells from 10 gonads) (Wilcoxon Tests). (E-G) Live imaging control (E) and Rho1-DN manipulated (F,G) ES15 gonads. Somatic F-actin, green. Cry2-Rho1DN-mCh, red. Yellow arrows, dispersed niche cells. (H) Quantified frequency of abnormal assembly with Rho1-DN in live analysis (p = 0.0003; Ctrl, n = 20 gonads; RhoDN, n = 19 gonads). (I-K) Immunostained gonads displaying niche assembly in ES17 gonads from photoactivated controls (I), dark state controls (J), and RhoDN-manipulated (K) gonads. Germ cells (Vasa, green), Niche (N-cad, magenta), and Nuclei (Hoechst, blue). (L) Quantified frequency of abnormal assembly with Rho1-DN in fixed analysis: Ctrl versus dark state (p = 0.6638; Ctrl, n = 36 gonads; Dark state, n = 22 gonads); Dark state versus RhoDN (p = 0.0017; Dark state, n = 22 gonads; RhoDN, n = 33 gonads); Ctrl versus RhoDN (p<0.0001; Ctrl, n = 36 gonads; RhoDN, n = 33 gonads) (Fisher’s Exact test). Time. HH:MM. Gonads and niches outlined. Scale bars, 10 um. n ≥ 3 trials. **Alt Text:** A-D show high zoom microscopy images of assembling niche cells with F-actin visualized in white. Graphs with statistical annotation quantifying the accumulation of F-actin at distinct cell interfaces at different developmental timepoints are shown below each image. E-G’’’ show microscopy images of control and manipulated gonads with F-actin and Cry2-mch labeled in distinct panels. H shows a graph with statistical annotation quantifying the frequency with which niches are abnormal in the earlier panels. I-K show fluorescent microscopy images of dissected and fixed tissue comparing control and optogenetically disrupted morphology, which is quantified in a statistically annotated graph in L.

Next to determine whether subcellular F-actin localization is required for niche assembly, we carried out persistent RhoDN recruitment through the assembly period with 488 nm blue light and 555 nm excitation to visualize mCh. As with OCRL live analysis experiments, manipulated animals inherited the *Six4*VP16-Gal4 genetic driver and had detectable cortical Cry2-RhoDN-mCh (Fig. 3F’,F’’’ and Fig. 3G’,G’’’), while sibling control animals did not inherit the driver and had no detectable Cry2-RhoDN-mCh (Fig. 3E’,E’’’). Manipulated gonads had visibly higher cortical relative to cytoplasmic Cry2-RhoDN-mCh after 4 hours of continued blue light exposure during live analysis (Compare 3F’&F’’’, 3G’&G’’’). We monitored niche assembly live in real time and analyzed the structure of end-stage niches. After four hours of blue light exposure, control gonads exhibited anterior assembly of niche SGPs (Fig. 3E’’). Live analysis with RhoDN recruitment showed abnormal niches that were either “misshapen” with irregular boundaries and not restricted to the anterior, or “dispersed” into multiple disconnected groups of cells (Fig. 3F’’,G’’). The misshapen niche phenotype supports the known requirement for Rho activity during compaction of the niche into its final architecture (Warder et al., 2024). Detection of dispersed niches phenocopies the defects seen with OCRL and reveals a role for Rho1-dependent F-actin localization in initial niche assembly. Abnormal niche morphology was detected significantly more often in manipulated gonads compared to sibling controls without the *Six4*VP16-Gal4 genetic driver (Fig. 3H).

We confirmed live RhoDN analysis with fixed tissue analysis after 6 hours of photoactivation. Fixed tissue experiments included light-exposed sibling controls and an additional dark-state control group. Dark-state controls were genetically identical to RhoDN manipulated animals but were never exposed to blue light. Light-exposed siblings controlled for general effects of blue light on tissue development, and the dark-state group controlled for effects from temporally regulated light-mediated RhoDN manipulation. RhoDN and dark-state control gonads had fewer germ cells than sibling controls without the driver, which is expected in animals from different genetic backgrounds (Compare Fig. 3I-K). Both sibling and dark-state control gonads had a single anterior niche with smooth boundaries, in contrast with RhoDN manipulated niches with irregular boundaries (Fig. 3I-K). Abnormal niche morphology was detected significantly more often in RhoDN gonads compared to both control groups (Fig. 3L), recapitulating the defects we detected with live analysis (Fig. 3H). While this result does not rule out the possibility of assembly contributions from adhesive sorting, it strongly supports a role for F-actin localization directing early niche assembly.

### Ena/VASP acts downstream of Rho1 to localize the niche F-actin cytoskeleton and mediate niche assembly

We asked which F-actin regulators Rho1 utilized to modulate cytoskeletal enrichment during niche assembly. The vasodilator-stimulator phosphoprotein (VASP), *Drosophila* Ena, is an anti-capping protein contributing to lamellipodial and filopodial protrusions (Krause et al., 2003). Ena reorganizes somatic F-actin to facilitate gonad coalescence earlier in development (Sano et al., 2012), and we hypothesized Ena might play a role in assembling niche cells.

We assessed subcellular Ena localization in assembling niches by immunostaining (Fig. 4A-D). We found Ena enriched at niche-GSC interfaces throughout assembly, with localization to the niche-GSC interface detected in gonads dissected at ES16 and becoming more robust at ES17 (Compare Fig. 5B and D). This enrichment precedes niche-GSC F-actin enrichment early in assembly and aligns with F-actin localization at assembly completion (Fig. 3A-B). With light-activated optogenetic RhoDN, the degree to which Ena was enriched at the niche-GSC interface was significantly reduced (Fig. 4E-H), suggesting Ena localization is affected by Rho function. This reduction occurred in both dark state and light activated gonads, suggesting that the minimal dark state interactions were sufficient to disrupt Ena enrichment. The disruption is further reduced in light activated gonads compared to dark state tissue, suggesting a temporal requirement for Rho function during niche assembly (Fig. 4D).

**Figure 4.**
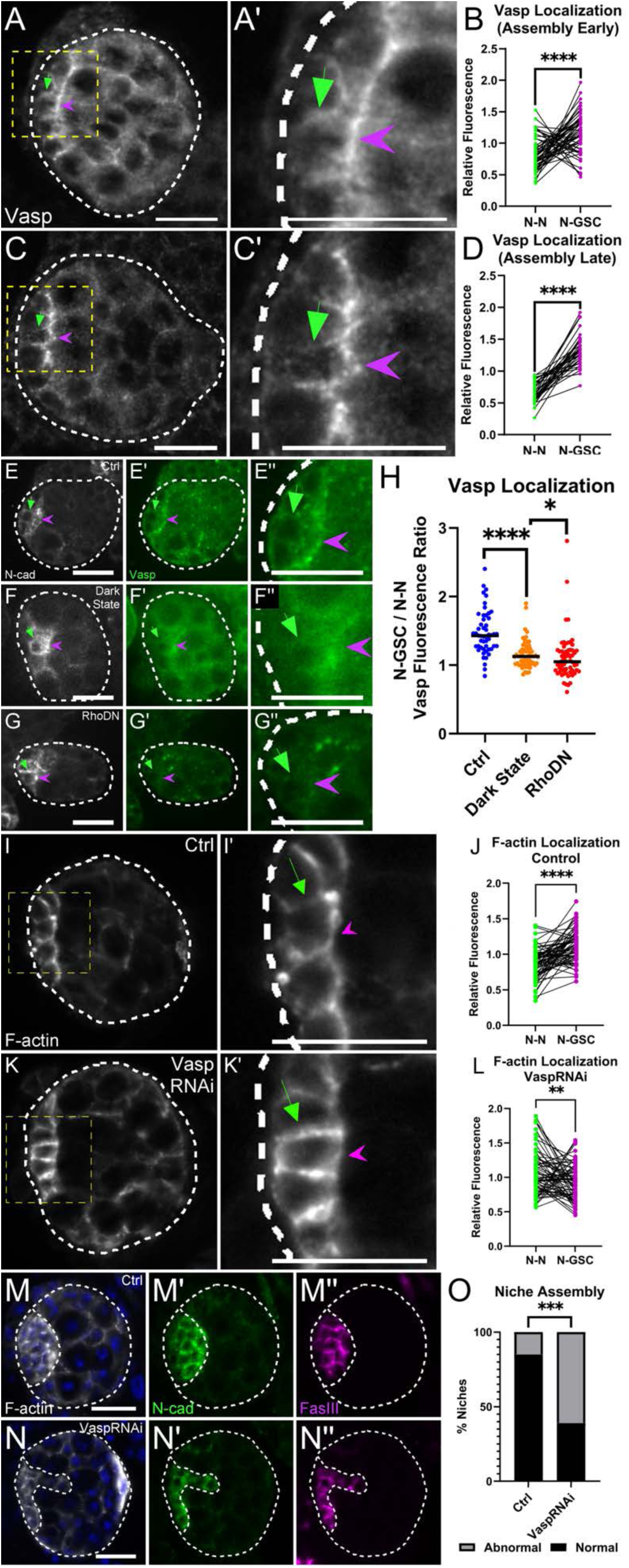
Rho1 mediates niche F-actin re-localization through Vasp. (A) ES16 and (C) ES17 dissected gonads showcasing Vasp localization (white) during early and late niche assembly. (B,D) Vasp enrichment at niche-niche (green) and niche-GSC (magenta) paired interfaces (B: p<0.0001; n = 54 from 17 gonads; D: p<0.0001; n = 37 from 12 gonads) (Mann-Whitney tests). (E) Photoactivated control, (F) dark state control, and (G) optogenetic RhoDN gonads, Niche (N-cad, white), Vasp (green). (H) Vasp enrichment at niche-GSC interface in photoactivated controls (blue), dark state controls (orange), and RhoDN manipulated gonads (red): Ctrl versus dark state (p<0.0001; Ctrl, n = 51 cells from 17 gonads; Dark state, n = 65 cells from 26 gonads); Dark state versus RhoDN (p = 0.0373; Dark state, n = 65 cells from 26 gonads; RhoDN, n = 61 cells from 24 gonads); Ctrl versus RhoDN (p<0.0001; Ctrl, n = 51 cells from 17 gonads; RhoDN, n = 61 cells from 24 gonads) (Mann-Whitney tests). (I,K) F-actin localization (white) in control and *vasp* RNAi gonads. (J,L) F-actin enrichment at niche-niche (green) and niche-GSC (magenta) paired interfaces (J: p<0.0001; L: p = 0.0033; each n = 67 cells from 22 gonads) (Wilcoxon tests). (M) Control and (N) *vasp* RNAi gonads (niche boundaries outlined: N-cad, green; Fas3, magenta). Somatic F-actin, white; Nuclei, Hoechst, blue. (O) Frequency of assembly defects with *vasp*RNAi (p = 0.0010; Ctrl, n = 26 gonads; *vasp*RNAi, n = 23 gonads) (Fisher’s Exact test). Niche-niche interfaces, green arrow; Niche-GSC interfaces, magenta arrowhead. Gonads outlined. Scale bars, 10 um. n ≥ 3 trials. **Alt Text:** Multiple sets of microscopy images show Vasp localization at subcellular interfaces in assembling niche cells, which is quantified with statistically annotated graphs. Additional sets of microscopy images show F-actin subcellular localization and niche morphology in Vasp disrupted gonads, accompanied by statistically annotated graphs that quantify the phenotypes displayed in the images.

**Figure 5.**
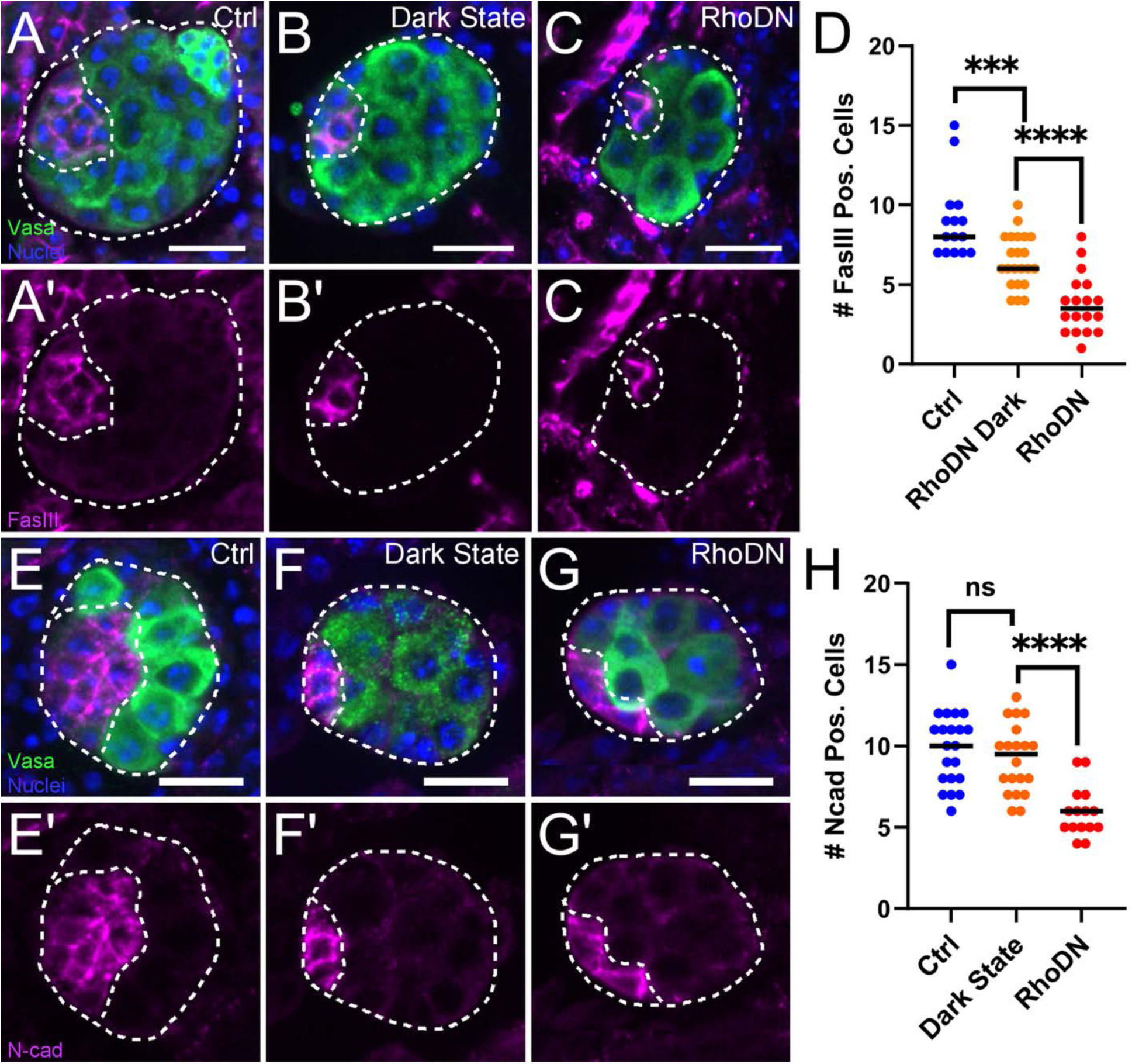
F-actin localization is required for the formation of niche cell contacts. (A-C) Single Z-slice images of gonads labeled with niche adhesion protein Fas3 (magenta). (D) Quantified niche cell counts in photoactivated controls (blue), dark state controls (orange), and RhoDN manipulated gonads (red): Ctrl versus dark state (p = 0.0008; Ctrl, n = 15 gonads; Dark state, n = 22 gonads); Dark state versus RhoDN (p <0.0001; Dark state, n = 22 gonads; RhoDN, n = 18 gonads); Ctrl versus RhoDN (p<0.0001; Ctrl, n = 15 gonads; RhoDN, n = 18 gonads) (Mann-Whitney tests). (E-G) Single Z-slice images of gonads labeled with niche adhesion protein N-cadherin (magenta). (H) Quantified niche cell counts: Ctrl versus dark state (p = 0.3777; Ctrl, n = 21 gonads; Dark state, n = 20 gonads); Dark state versus RhoDN (p <0.0001; Dark state, n = 20 gonads; RhoDN, n = 15 gonads); Ctrl versus RhoDN (p<0.0001; Ctrl, n = 21 gonads; RhoDN, n = 15 gonads) (Mann-Whitney tests). Germ cells (Vasa, green) and Nuclei (Hoechst, blue). Scale bars, 10 um. n ≥ 3 trials. **Alt Text:** Microscopy images of dissected gonads labeled to show germ cells, nuclei, and niche-specific adhesion proteins Fasciclin3 or N-cadherin in magenta. Images represent a single Z slice. Graphs show quantification of the number of all niche cells per gonad for several gonads, plotted for light-exposed controls, dark-state, and RhoDN tissue. Graphs are statistically annotated.

We evaluated a role for Ena in niche cytoskeletal localization and niche assembly with RNAi knockdown in SGPs, assessing each with fixed analysis at ES17 when niche assembly should have completed. Controls exhibited F-actin enrichment at niche-GSC interfaces (Fig. 4I-J). With *ena* knockdown, F-actin instead localized to niche-niche interfaces (Fig. 4K-L). These results suggest that while Ena is not required for initial F-actin localization at niche-niche interfaces, it is required for later relocalization to the niche-GSC interface. This role is consistent with the enrichment of Ena at the niche-GSC interface prior to F-actin relocalization there (Fig 4A-B). The localization of F-actin to the incorrect interface with *ena* reduction differs from the total loss of local enrichment seen with RhoDN (Fig. 3C-D), suggesting that *ena* is not the only Rho effector modulating F-actin localization. SGP *ena* knockdown disrupted niche assembly, with niches frequently exhibiting irregular boundaries and dispersed cells as detected with optogenetic RhoDN (Fig. 4I-K). Together, these results suggest that Rho affects Ena enrichment at the niche-GSC interface, where Ena modulates F-actin localization to impact niche architecture.

### Localized F-actin is necessary to form niche cell contacts and manifest identity during assembly

Since F-actin modulators regulate trafficking of adhesion proteins to specific cell interfaces (Cordova-Burgos et al., 2021), we asked if F-actin localization was required for establishment of adhesion between niche cells. If F-actin localization were required to establish adhesive contacts, then niche cells would be unable to undergo adhesive sorting in the absence of F-actin regulation. We assessed Fasciclin3 (Fas3) and N-cadherin in RhoDN photoactivated animals compared to both dark state and light-exposed controls that lacked the genetic driver.

Light-activated RhoDN gonads had significantly fewer niche cells accumulating either Fas3 or N-cadherin (Fig. 5). Dark state controls had fewer Fas3 cells than light-exposed controls, consistent with the low levels of dark state activity we noted with this system (Fig. 1I-K). The number of Fas3+ niche cells was further reduced upon light activation, supporting the temporal requirement for Rho during assembly (Fig. 5D). The number of N-cad positive niche cells was significantly reduced in RhoDN manipulated animals, but unaffected in dark state tissue compared to light activated controls (Fig. 5H). These results suggest F-actin localization is required for accumulation of niche adhesion proteins. Adhesive sorting would not be possible without accumulation of adhesion proteins. Thus, these results suggest that F-actin localization is required to enable adhesive interactions among niche cells that contribute to initial niche assembly.

Our previous work showed that niche identity defects could lead to loss of niche adhesion (Anllo and DiNardo, 2022; Hofe et al., 2026). Thus, we tested whether niche specification is affected in RhoDN experiments. We assessed Hedgehog (Hh) and Islet accumulation, which are enriched in properly specified niche cells (Anllo and DiNardo, 2022; Hofe et al., 2026; Michel et al., 2012). When F-actin localization was disrupted with light-activated RhoDN, niche cells accumulated significantly less Hh and Islet (Fig. 6). We did not detect a reduction in either identity marker when comparing dark state to light-exposed controls, indicating that identity defects resulted from temporal, light-mediated Rho disruption during assembly (Fig. 6D,H). These results suggest defects in niche identity when F-actin localization is lost during assembly.

**Figure 6.**
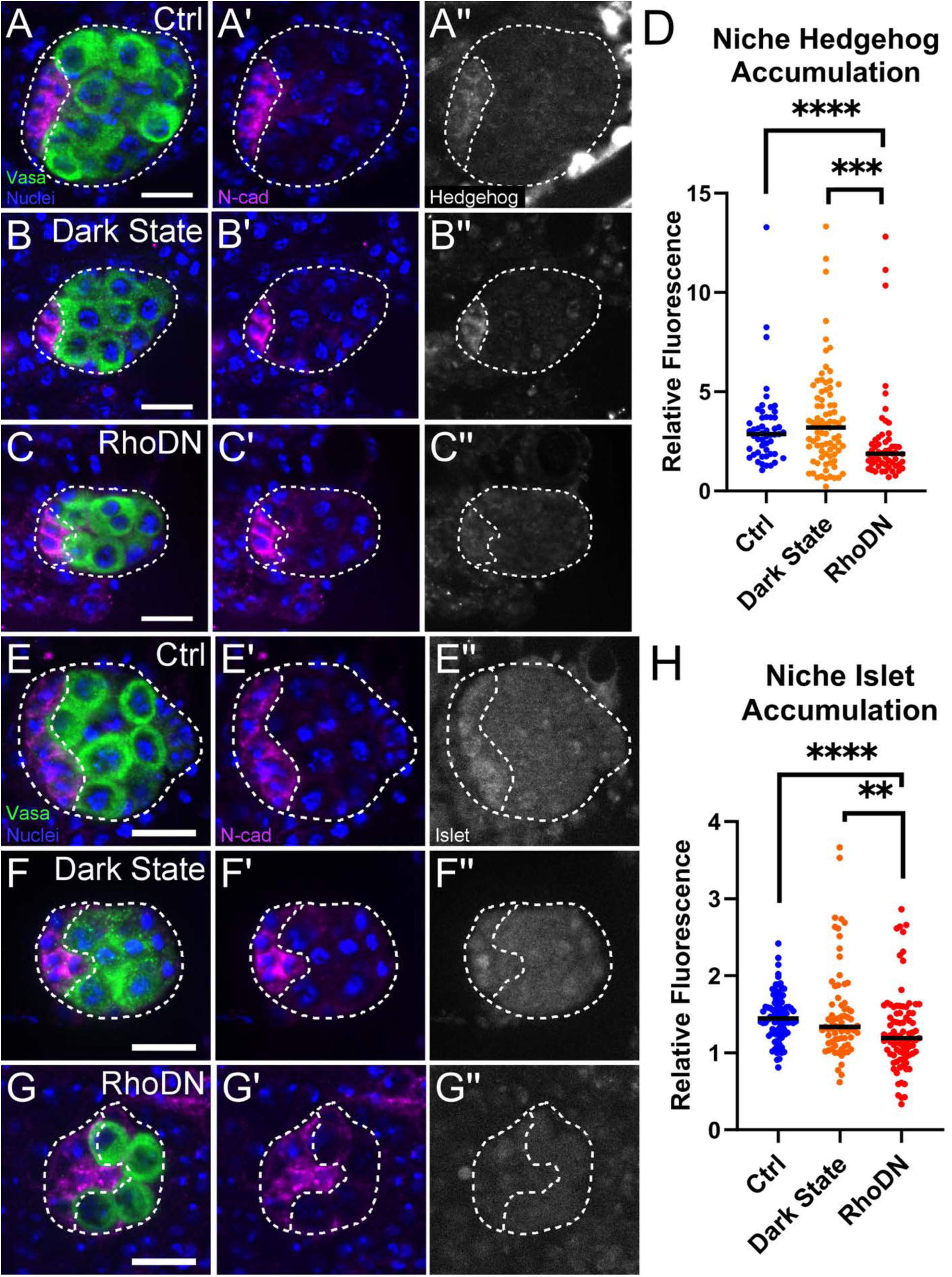
F-actin localization during niche assembly is required for the maintenance of niche identity. (A-C) Hedgehog (white) in niche cells (N-cad, magenta). (D) Quantified Hedgehog accumulation within niche cells in photoactivated controls (blue), dark state controls (orange), and RhoDN manipulated gonads (red): Ctrl versus dark state (p = 0.5073; Ctrl, n = 51 cells from 17 gonads; Dark state, n = 81 cells from 26 gonads); Dark state versus RhoDN (p = 0.0003; Dark state, n = 81 cells from 26 gonads; RhoDN, n = 57 cells from 20 gonads); Ctrl versus RhoDN (p<0.0001; Ctrl, n = 51 cells from 17 gonads; RhoDN, n = 57 cells from 20 gonads) (Mann-Whitney Tests). (E-G) Islet (white) in niche cells (N-cad, magenta). (H) Quantified Islet accumulation within niche cells: Ctrl versus dark state (p = 0.2798; Ctrl, n = 93 cells from 31 gonads; Dark state, n = 75 cells from 25 gonads); Dark state versus RhoDN (p = 0.0076; Dark state, n = 75 cells from 25 gonads; RhoDN, n = 87 cells from 29 gonads); Ctrl versus RhoDN (p<0.0001; Ctrl, n = 93 cells from 31 gonads; RhoDN, n = 87 cells from 29 gonads) (Mann-Whitney Tests). Germ cells (Vasa, green) and Nuclei (Hoechst, blue). Scale bars, 10 um. n ≥ 3 trials. **Alt Text:** Microscopy images of dissected gonads labeled to show germ cells (green), nuclei (blue), and niche adhesion protein N-cadherin (magenta) and either Hedgehog or Islet markers of niche-specific gene expression in white. Graphs show quantified fluorescence intensity of either Hedgehog or Islet in the niche plotted for light-exposed controls, dark-state, and RhoDN tissue. Graphs are statistically annotated.

Niche specification requires early Notch signaling at 5-6 hours after egg laying (Okegbe and DiNardo, 2011), but evidence of niche-specific gene expression is not visible until niche assembly after pro-niche SGPs make contact (Anllo et al., 2019; Le Bras and Van Doren, 2006). Since embryos were light-activated after the early requirement for Notch activity, the manipulations to F-actin localization are occurring after this signaling event. Thus, F-actin localization during niche assembly is important for SGPs to attain niche identity after the initial signaling required for assignment of niche fate earlier in development.

### Polarized F-actin is required for niche function

Since proper niche architecture and identity are required for function (Anllo and DiNardo, 2022; Vida et al., 2025; Warder et al., 2024), we asked how F-actin localization manipulations affected the ability of the niche to support resident stem cells. Bona fide niche cells signal to GSCs, resulting in accumulation of the protein STAT. In the adult, loss of STAT leads to GSC abscission failure during division, and failure to release GSC daughter cells from the niche to enable differentiation (Lenhart and DiNardo, 2015). Upon disrupting F-actin localization with RhoDN, we detected reduced GSC STAT accumulation, with no detectable effect in dark state tissue (Fig. 7A-D). Niche signaling also biases the orientation of cell division perpendicular to the niche-facing germ cell cortex, with one centrosome located at this cortex. This orientation permits one cell to remain in the niche with stem identity and the other to be displaced and differentiate (Chen et al., 2018; Kiger et al., 2001; Leatherman and DiNardo, 2008; Tulina and Matunis, 2001; Yamashita et al., 2003). We detected proper centrosome alignment with the niche-GSC interface in light-exposed and dark state controls (Fig. 7E-F, H). However, light mediated RhoDN disrupted centrosome localization, resulting in both centrosomes occasionally positioned away from the niche interface (Fig. 7G,H). Because we found an effect of Rho on Ena enrichment, we also assessed STAT accumulation and centrosome localization with *ena* knockdown and found both defective (Fig. 8). Thus, F-actin localization disruptions impair niche cell signaling to stem cells and proper division control. These functionality defects are consistent with expectations for compromised niche identity.

**Figure 7:**
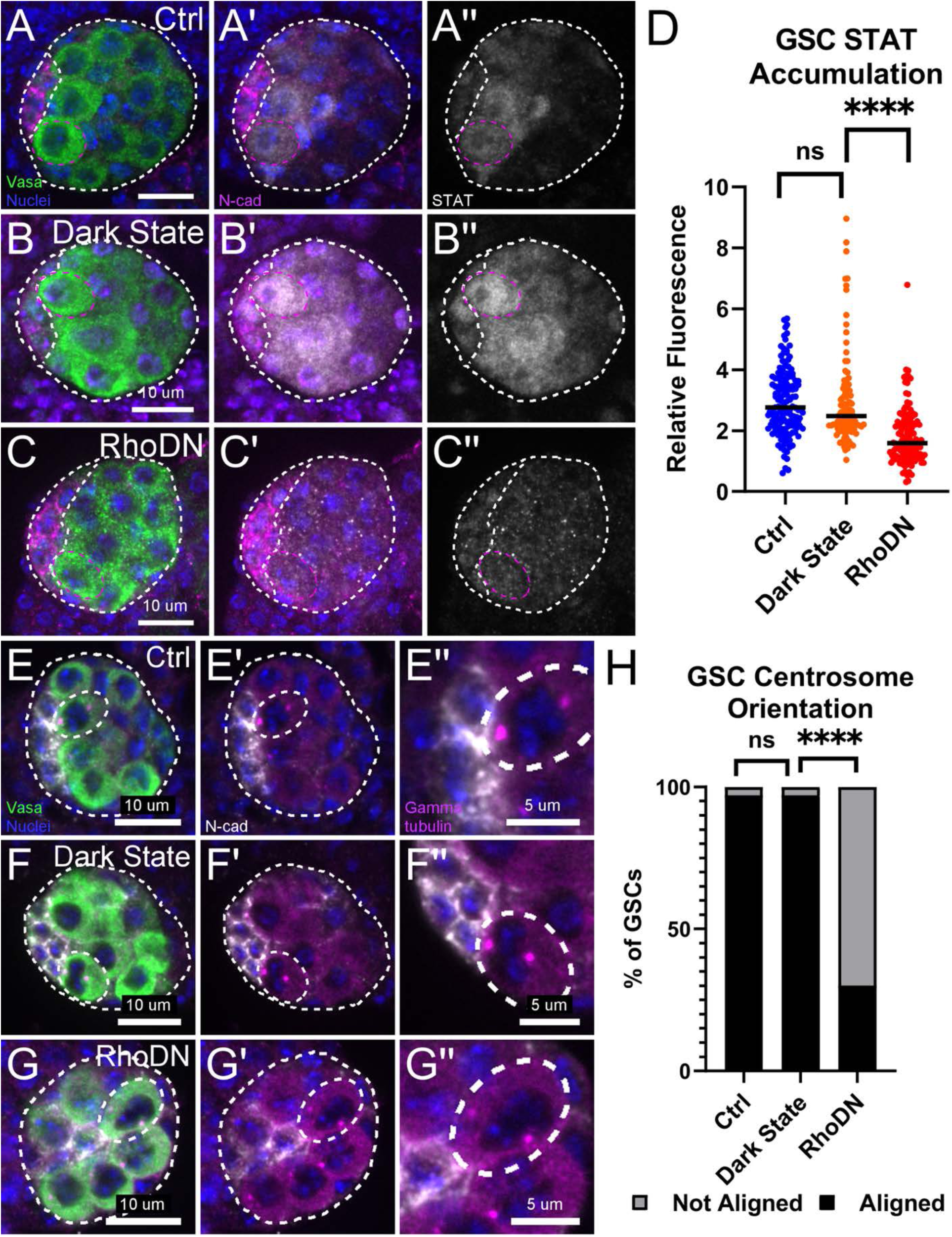
Localized F-actin is necessary to form a functional niche. Fixed analysis of photoactivated controls (Ctrl), dark state controls (Dark State), and optogenetic RhoDN manipulated (RhoDN) gonads. (A-C) Stat accumulation (white) in GSCs (magenta outline). Niche cells, magenta (N-cad). (D) Quantification of GSC Stat in photoactivated controls (blue), dark state controls (orange), and RhoDN manipulated gonads (red): Ctrl versus dark state (p = 0.2798; Ctrl, n = 145 GSCs from 29 gonads; Dark state, n = 95 from 19 gonads); Dark state versus RhoDN (p<0.0001; Dark state, n = 95 from 19 gonads; RhoDN, n = 125 from 25 gonads); Ctrl versus RhoDN (p<0.0001; Ctrl, n = 145 GSCs from 29 gonads; RhoDN, n = 125 GSCS from 25 gonads) (Mann-Whitney tests). (E-G) Centrosomes (Gamma tubulin, magenta) and niche cells (N-cad, white). Dotted line shows GSCs with two centrosomes. (H) Quantified centrosome orientation in GSCs with two centrosomes where the centrosomes are aligned (black) or not aligned (grey): Ctrl versus dark state (p>0.9999; Ctrl, n = 97 GSCs from 32 gonads; Dark state, n = 55 from 19 gonads); Dark state versus RhoDN (p<0.0001; Dark state, n = 95 GSCs from 19 gonads; RhoDN, n = 81 from 24 gonads); Ctrl versus RhoDN (p<0.0001; Ctrl n = 97 GSCs from 32 gonads; RhoDN, n = 81 from 24 gonads) (Fisher’s exact tests). Germ cells (Vasa, green) and Nuclei (Hoechst, blue). n ≥ 3 trials. **Alt Text:** A-C show microscopy images of dissected gonads immunostained to show STAT accumulation in germline stem cells. D shows a statistically annotated graph comparing STAT accumulation in the images. E-G show microscopy images of dissected gonads immunostained to show centrosome localization, which is quantified in a statistically annotated graph in H.

**Figure 8:**
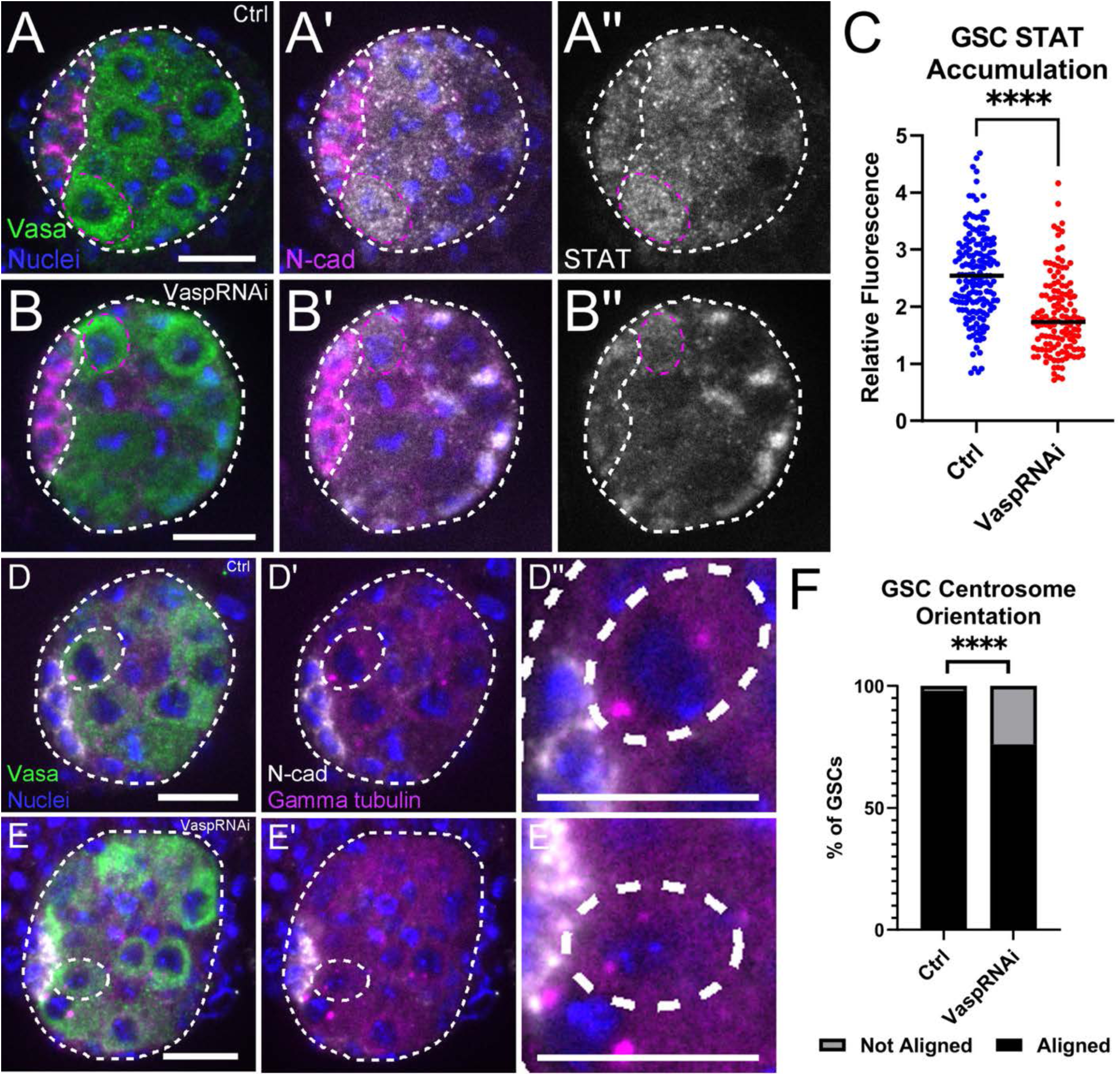
Vasp localizes niche cell F-actin required for functionality. (A) Control and (B) *vasp* RNAi gonads immunostained for Stat accumulation (white) in GSCs (magenta outline). Niche, outline, N-cad, magenta. (C) Reduced GSC Stat in *vasp*RNAi (red) versus control (blue) (p<0.0001; Ctrl, n = 160 GSCs from 32 gonads; *vasp* RNAi, n = 130 from 26 gonads) (Mann-Whitney). (D) Control and (E) *vasp* RNAi gonads immunostained for centrosomes (Gamma tubulin, magenta, white outline). Germ cells, Vasa, green; Niche, N-cad, white; Nuclei, Hoescht, blue. (F) Quantified centrosome orientation in GSCs with two centrosomes (p<0.0001; Ctrl n = 65 GSCs from 29 gonads; *vasp* RNAi, n = 104 from 31 gonads) (Fisher’s exact test). Germ cells (Vasa, green) and Nuclei (Hoechst, blue). n ≥ 3 trials. **Alt Text:** A-B show microscopy images of dissected gonads immunostained to show STAT accumulation in germline stem cells. C shows a statistically annotated graph comparing STAT accumulation in the images. D-E show microscopy images of dissected gonads immunostained to show centrosome localization, which is quantified in a statistically annotated graph in F.

## Discussion

Our results reveal a direct requirement for F-actin localization during assembly for niche cells to properly undergo morphogenesis and manifest their cellular identity, which is subsequently crucial for niche tissue function. We show that niche F-actin localization depends on somatic Rho1 during assembly, and is required for niche cells to accumulate adhesion proteins and establish niche specific gene expression. We identify *Drosophila* Ena/VASP as one cytoskeletal modulator whose accumulation is impacted by Rho1 and relevant for niche morphology and function. This work also demonstrates the Cry2-CIB system as a method for tissue-specific cytoskeletal manipulation with high temporal precision during gonadogenesis, presenting opportunities to define mechanisms underlying gonadal cell behaviors at precise developmental timepoints.

### F-actin localization directly enables niche assembly

Our previous work identified F-actin enrichment at precise niche cell interfaces during assembly (Anllo and DiNardo, 2022; Warder et al., 2024). Localized F-actin can drive morphological changes that enable tissue formation. Alternatively, establishment of adhesive cell contacts could localize cytoskeletal regulators in response to morphogenesis (Tsai et al., 2022). For example, during neural crest migration, a switch from E-cadherin to N-cadherin drives a shift in force from cell-cell contacts to free cell edges, resulting in dispersal of migratory neural crest cells and change in localization of F-actin regulators (Scarpa et al., 2015). Upon disrupting F-actin localization during assembly, we find fewer niche cells establishing adhesive contacts. This effect is not a simple result of failed motility, as genetic disruptions to niche assembly signals do not impact the number of adhesive niche cells. While these disruptions ultimately lead to dispersed, non-anterior niche cell associations, loss of motility cues does not reduce the number of niche cells accumulating Fas3 or E-cadherin (Anllo and DiNardo, 2022). Our results support a specific role for F-actin localization during the developmental timepoint of niche assembly controlling adhesive protein accumulation in the forming niche. Thus, F-actin localization in pro-niche cells during assembly ultimately drives morphogenesis impacting cell contact formation and tissue assembly.

### Localized F-actin during assembly is required to establish niche identity

Niche identity assignment requires early SGP receipt of Notch signaling. Previous studies found Notch activation at embryonic stages 11-12 was sufficient to enable niche specification, hours before gonad coalescence and niche assembly (Okegbe and DiNardo, 2011). Notch activation has been detected broadly in SGPs throughout the gonad prior to niche establishment (Kitadate and Kobayashi, 2010; Okegbe and DiNardo, 2011). Early and unlocalized activity suggests Notch signaling renders SGPs competent to attain niche identity. Our optogenetic manipulations are precisely timed to introduce blue light only after the early developmental requirement for Notch activation and during niche assembly, when SGPs have already been rendered competent to assume niche fate. The reduced Islet and Hedgehog accumulation and disrupted communication with GSCs suggests failure to assume bona fide niche identity when F-actin localization is perturbed during this later developmental timepoint. These data suggest localized F-actin establishing pro-niche cell contacts is a requirement to enable niche-specific gene expression following earlier competence rendered by Notch.

In addition to Notch activation, repressive EGFR signaling from posterior germ cells is required to restrict niche cell fate to anterior SGPs. Germ cell reduction results in specification of additional niche cells (Kitadate and Kobayashi, 2010). Of note, gonads dissected from the genetic background that yielded dark state and optogenetically manipulated tissue had relatively few germ cells compared to other genetic backgrounds. With fewer germ cells, we would anticipate formation of supernumerary niche cells. It is thus possible that the reduced niche cell number we detect with Rho disruption is even less severe than it would be in a genetic background with additional germ cells as sources for repressive niche identity cues.

Niche tissue function is dependent on its ability to send self-renewal signals and stem cell division orientation cues (Leatherman and DiNardo, 2010; Tulina and Matunis, 2001; Yamashita et al., 2003). We find reduced accumulation of STAT in GSCs, and misorientation of GSC centrosomes upon disruption of F-actin localization, suggesting that both receipt of niche signals and direction of cell division is impaired. This lack of functionality is likely a direct result of defective niche cell identity.

Our data support a role for F-actin directed establishment of niche cell contacts in manifestation of niche identity. In other genetic disruptions to signals required for niche assembly, niche cells are dispersed and do not reach the anterior, but the number of cells that accumulate niche adhesion proteins is unchanged and these cells retain expression of the niche identity gene *upd* (Anllo and DiNardo, 2022). Thus, disruptions to niche cell motility do not interfere with niche identity per se. The loss of niche identity we detect is thus not simply an outcome of failed migration. Our finding that niche F-actin localization is required for identity is an exciting concept that could not have been directly inferred from previous work in this system.

### Rho1 mediates establishment of a functional niche through factors including VASP

Our work establishes a role for Rho activity in localizing F-actin to enable assembly of the testis niche. Previous studies have shown a role for Rho kinase, an important downstream Rho effector, in defining and maintaining proper niche shape after cell assembly, during tissue compaction and steady state adult function. These studies utilized different genetic drivers to enable disruption later in development or only in adulthood. During these later stages, disruptions to Rho kinase activity caused defects in niche architecture but did not impact niche cell number or identity (Vida et al., 2025; Warder et al., 2024). The misshapen niches resulting from Rho kinase disruptions after assembly caused niche contact with additional germ cells, with significantly more STAT+ GSCs (Warder et al., 2024). Niche cells retained expression of the identity gene *upd* when Rho kinase was compromised during adulthood (Vida et al., 2025). These defects contrast with those we detect when disrupting Rho earlier in development, when niche cells are in the process of migrating to their final assembly point at the embryonic gonad anterior. Earlier disruptions in the present study result in specification of fewer niche cells with reduced expression of niche identity markers and failure to accumulate STAT in adjacent germ cells (i.e. failed induction of GSC identity). Collectively, our results and published studies indicate a role for Rho activity in contributing to niche identity during assembly, with that role changing to one required for maintenance of niche architecture later in development.

A direct role for Rho1 specifying niche cells is unlikely, as known cytoskeletal influences on gene expression require force transduction (Park et al., 2023). Alternatively, F-actin localization might influence niche identity by promoting cell contacts that enable signaling between adjacent cells. We detect fewer Fas3 and Ncad positive niche adhesions in the absence of F-actin localization, which could reflect lack of architectural features required to transmit forces or facilitate signaling at forming niche-niche interfaces.

We find mislocalization of Ena/Vasp with disrupted Rho. While overall levels of Vasp protein are unaffected, the degree to which Vasp accumulates at niche-stem cell interfaces relative to niche-niche interfaces is significantly reduced. Likewise, Vasp disruption results in failed relocalization of F-actin to the niche stem cell interface after assembly, and compromised niche signaling to stem cells. While our results support an effect of Rho on Vasp, Vasp disruption does not result in complete loss of F-actin polarity as seen with disrupted Rho. We expect that Vasp is one of multiple relevant downstream targets required to polarize the niche F-actin cytoskeleton during initial niche assembly.

### Cry2-CIB optogenetics can interrogate morphogenesis of internally positioned tissues with temporal precision

Genetic drivers allow dissection of tissue-specific contributions to morphogenesis but are often expressed during broad developmental timepoints. Previous studies of cytoskeletal contributions to niche morphogenesis were complicated by earlier requirements for somatic cytoskeletal regulation enabling gonad coalescence, and later roles for F-actin contractility in compacting the assembled niche (Sano et al., 2012; Warder et al., 2024). Optogenetic control of cytoskeletal regulators overcomes this obstacle, allowing temporal disruption precisely during the developmental timeframe when pro-niche cells are migrating to their anterior assembly point. Previous optogenetic cytoskeletal disruptions were performed in the developing *Drosophila* epidermis, a peripheral tissue positioned more accessibly for light manipulation (Guglielmi and De Renzis, 2017; Guo et al., 2022). The embryonic *Drosophila* gonad is positioned up to 100 microns more internal than the epidermis, complicating accessibility for optogenetic control. Our validation of the ability to mobilize subcellular protein localization, disrupt cytoskeletal localization, and affect tissue morphogenesis is an important technical innovation enabling precise temporal manipulation of internally developing tissues. In combination with tissue-specific drivers, this advance enables incisive assessment of cytoskeletal contributions to specific stages of morphogenesis during embryogenesis.

## Data availability

The *Drosophila melanogaster* strains used in this work are available either publicly or upon request. All relevant data and details of resources can be found within the article, figures, tables, and supplementary information.

## Supporting information

Supplemental Table 1 - Resource Table

Supplemental Table 2 - Genotype Table

## Acknowledgements

We thank K. Lenhart, S. DiNardo, and E.T. Ables for manuscript comments, the Bloomington *Drosophila* Stock Center (NIH P40OD018537), R. Lehmann, B. Warder, S. DeRenzis, and B. He for stocks, and E. Bach for antibodies.

